# Predicting the biological invasion risks of the most farmed insect for food and feed

**DOI:** 10.1101/2025.10.30.685499

**Authors:** Eléna Manfrini, Franck Courchamp, Pierre-Olivier Maquart, Quentin Lamboley, Boris Leroy

## Abstract

Insect farming is recognized as a sustainable alternative to traditional livestock, offering significant reductions in land, water, and feed use, along with lower greenhouse gas emissions. However, the potential for farmed insects to escape or be released poses a significant biological invasion risk, threatening biodiversity, human well-being, and economies. This study is the first to quantify the global spatial invasion risk of seven of the most farmed insect species. We employ state-of-the-art Species Distribution Models integrating climatic and socio-economic variables to predict habitat suitability and identify high-risk regions. Our analysis highlights substantial invasion risks globally, with key hotspots in the Caribbean, South America, Western and Central Europe, and Oceania. While high-risk areas are species-dependent, the banded cricket, common housefly, and black soldier fly pose the greatest risks. Notably, nearly all surveyed insect farms are located in areas with intermediate to high invasion risks, underscoring the urgent need for stringent biosecurity measures in existing farms and strategic species selection for future farms. By providing a robust methodology and guidelines, our work offers a foundation for policymakers and industry stakeholders to ensure sustainable insect farming and critical insights into this understudied field.

## Introduction

The global human population is projected to reach 10.3 billion by mid 2080s^1^, driving a 35 to 56% increase in food demand as soon as 2050^2^. Meeting this demand sustainably is a critical challenge, as conventional food production—particularly livestock farming—drives land-use change, greenhouse gas emissions, and biodiversity loss^3,4^, all of which impact biodiversity and nature’s contributions to people^5^. Ensuring food security while minimizing environmental impacts requires innovative solutions that align with the potentially conflicting United Nation sustainability goals of *Zero hunger* (2) vs *Climate action* (13) and *Life on land* (16)^6^.

Insect farming has emerged as a promising alternative to traditional livestock production, requiring less land, water, and feed (some species even eat organic waste), and emitting significantly fewer greenhouse gases and ammonia^7,8^. It could also support food security by providing affordable protein, especially where meat is scarce or expensive^9^. Insect-based products are increasingly accepted for human consumption and are already widely used for aquaculture and animal feed^7^. In Europe, eight insect species are approved for human food and animal feed^10,11^: the black soldier fly (*Hermetia illucens*), the yellow mealworm (*Tenebrio molitor*), the house cricket (*Acheta domesticus*), the lesser mealworm (*Alphitobius diaperinus*), the common housefly (*Musca domestica*), the banded cricket (*Gryllodes sigillatus*), the field cricket (*Gryllus assimilis*) and the migratory locust (*Locusta migratoria*). However, insect farming also presents unresolved risks to biodiversity and human society^12^. Notably, if farmed outside their native ranges, insects can pose a significant biological invasion risk^13^.

Biological invasions are defined by the human introduction, establishment, and successful spread of species populations outside their natural range^14^. These invasions are implicated in over 65% of documented species extinctions and cost over $2.168 trillion to global economies^15^, which is as much as natural hazards^16^. Insects are among the most invasive taxonomic groups, with economic impacts exceeding US $5 trillion from 1960 to 2020^17,18^, in addition to significant ecological impacts^19^. Diverse examples are agricultural losses, like the fall armyworm *(Spodoptera frugiperda*) in Africa, Asia and Oceania^20^, health issues from diseases spread by tiger mosquitoes (*Aedes aegypti* and *Ae. albopictus*)^21,22^, and ecosystem disruptions by the red imported fire ant (*Solenopsis invicta*), which reduces native invertebrate and vertebrate populations^23^.

After 1950, the globalization of trade has exponentially increased these risks, with species introductions from all continents into all continents through various pathways^24–26^, including animal farming as a primary pathway of invasion^27,28^. Introductions due to animal farming, either intentional (e.g., translocations) or non intentional (e.g., insufficient biosecurity measures) have been almost always systematic^14,28^. Large-scale rearing facilities can produce millions of individuals per week, with the total feed produced worldwide in 2011 amounting approximately to 870 million tons^9^ and is still increasing globally^29–31^.

The best way to prevent biological invasions is to avoid introducing species into suitable ecosystems outside their native range^32^. Because of the almost systematic nature of escapees from facilities in animal farming, this implies not rearing species in regions where, if released or escaped, they could establish in the wild. To estimate suitable habitats of non-native species, and therefore their risk of establishment, spread and invasion, one can rely on Species Distribution Models (SDMs) which statistically estimate the relationship between a species and its environmental, bioclimatic, and socio-economic drivers^33^. Our objective with this study is to resolve the question of the spatial invasion risks posed by large-scale insect farming, using SDMs. We focused on the eight aforementioned species, which are among the most studied and farmed species globally^34^. We address this question by first projecting the spatial risks of invasions, and then comparing this spatial risk with known locations of existing farming sites.

## Results

### Model performance and selections for global predictions

We evaluated several climatic and socio-economic variables as potential predictors of species occurrences. Our variable selection process retained four key climatic variables: bio5, bio6, hurs_min, and npp (**Table 1**) and one socio-economic variable (croplands). Among the socio-economic variables, we excluded the *Human population* variable due to its minimal influence on the models, and the *Human footprint* variable because of its high correlation with *Croplands* (**Supplementary Fig. S1**). In addition, we retained both *bio5* and *bio6* despite their correlation, as our evaluation process revealed that using only one of these two variables reduced the ecological realism of species-environment relationships (**Supplementary Fig. S2)**.

**Table 1:**
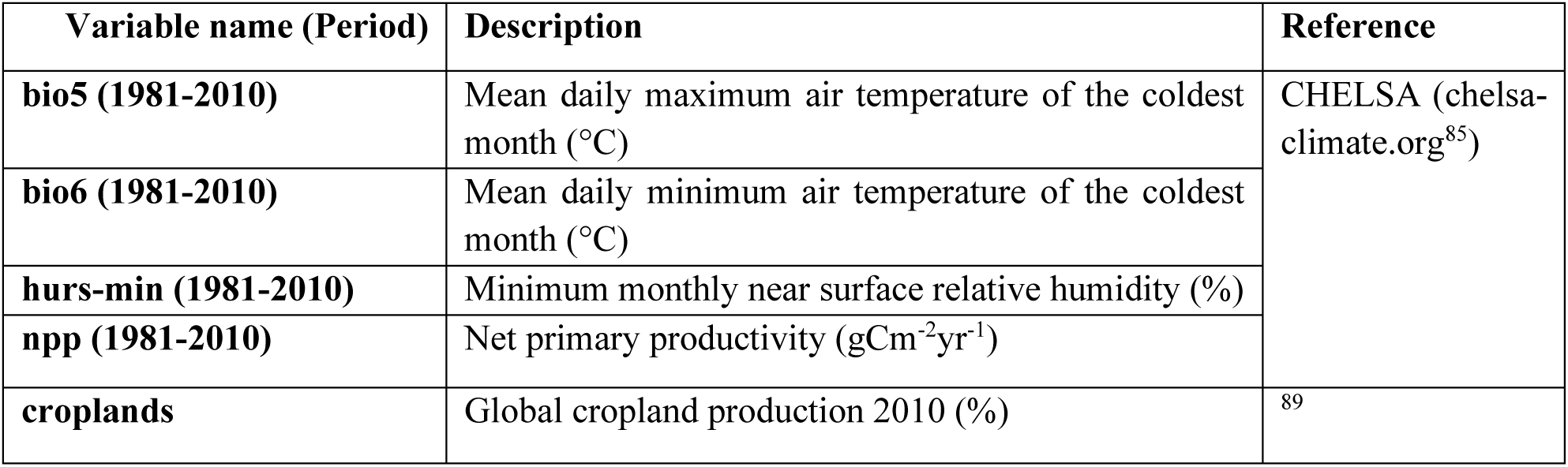
Final environmental predictors used to model species distributions.

| **Variable name (Period)** | **Description** | **Reference** |
| --- | --- | --- |
| **bio5 (1981-2010)** | Mean daily maximum air temperature of the coldest month (°C) | CHELSA (chelsa-climate.org<sup>85</sup>) |
| **bio6 (1981-2010)** | Mean daily minimum air temperature of the coldest month (°C) |  |
| **hurs-min (1981-2010)** | Minimum monthly near surface relative humidity (%) |  |
| **npp (1981-2010)** | Net primary productivity (gCm<sup>-2</sup>yr<sup>-1</sup>) |  |
| **croplands** | Global cropland production 2010 (%) | <sup>89</sup> |

Among the five modelling techniques tested (MaxNet, Extreme Gradient Boosting XGBOOST, down-sampled random forest RF, generalized additive models GAM, generalized linear models GLM), only MaxNet, RF and GLM succeeded in our evaluation criteria (Boyce Index > 0.7) for all but one species (**Supplementary Fig. S3**). For this species, the field cricket, nearly half of the models did not perform adequately (**Supplementary Fig. S3**) and selected models produced ecologically unrealistic response curves (**Supplementary Fig. S4**). Consequently, we excluded this species from further analysis. For the remaining seven species, XGBOOST and GAM models frequently did not meet performance criteria, with none of the GAM models being selected for two species (common housefly and migratory locust, **Supplementary Fig. S3**). Additionally, some selected GAM and XGBOOST models produced ecologically unrealistic response curves, including a total absence of response for some variables (**Supplementary Fig. S4**). Therefore, our final ensemble model was based on the average suitability value among MaxNet, RF and GLM models.

### Suitability predictions and spatial invasion risk of the seven most farmed insect species

The spatial habitat suitability projections for the seven most farmed species indicate a wide range of potential establishment sites worldwide (**Fig. 1**; see suitability and standard deviation maps for all species in **Supplementary Fig. S5**), thereby posing invasion risks (**Supplementary Fig. S6**). Specifically, these species exhibit a high risk of invasion in regions such as eastern Canada, the western United States, Uruguay, Europe, South Africa, the East Asian coast, Southeast Australia, and New Zealand (**Fig. 2**). Conversely, the invasion risk appears lower in the northern regions of the Northern Hemisphere. Our model projections identified two species with large areas of high invasion risk (black soldier fly & banded cricket), as well as a third species with large areas of intermediate invasion risk (housefly) (**Fig. 3**). The four other species were projected to be able to invade a more limited number of countries with a high risk of establishment (around 17 countries on average), yet many countries with an intermediate risk of establishment (44).

**Figure 1:**
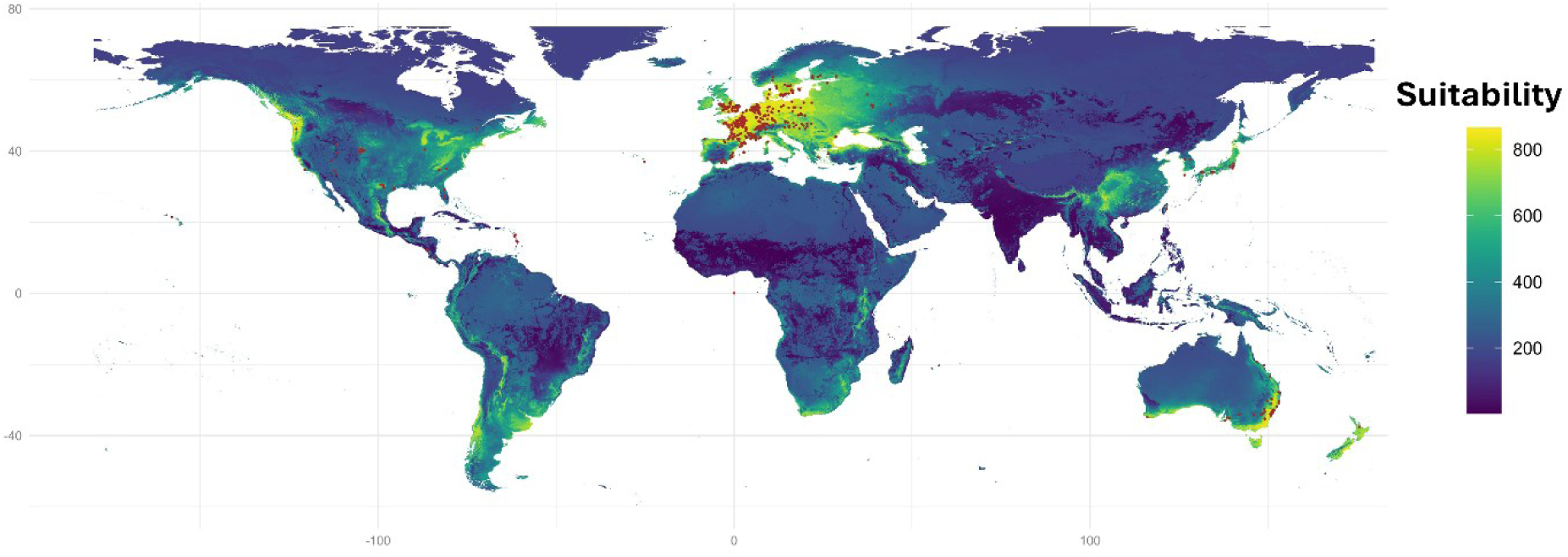
Species predicted suitability maps of the lesser mealworm. See these maps for all species in **Supplementary Figure S5,A**. The suitability map displays species occurrences as red points.

**Figure 2:**
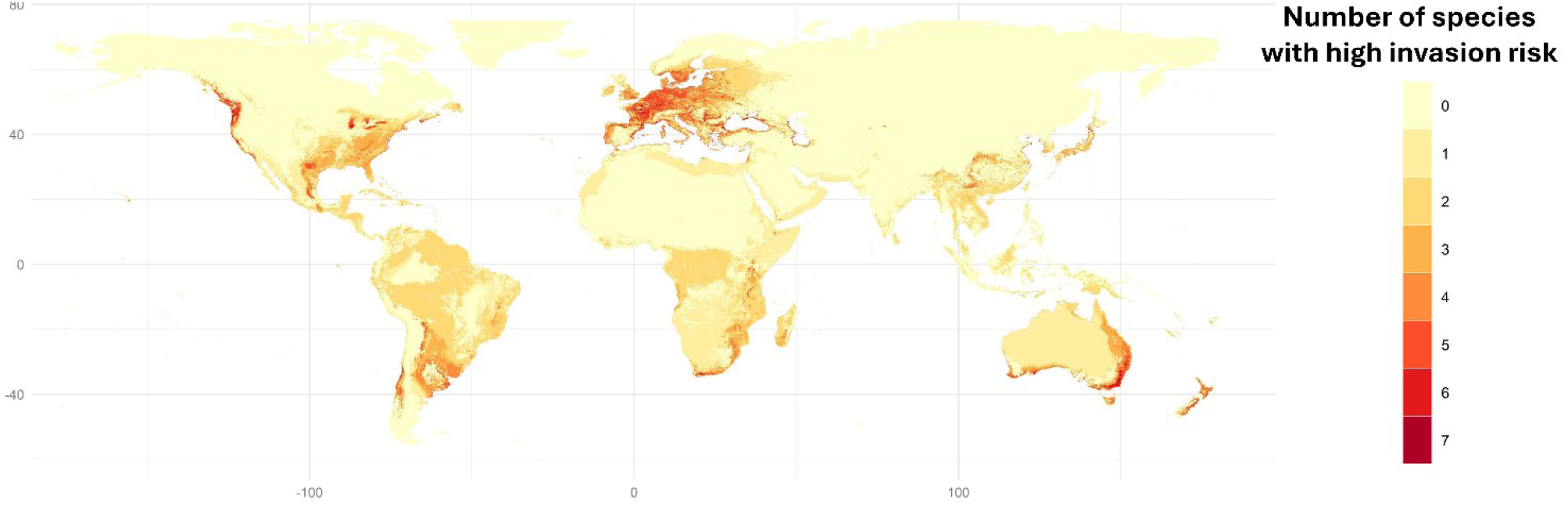
Global distribution of high invasion risk, cumulated for the seven most farmed insect species. This map illustrates the number of species per pixel exhibiting high invasion risk.

**Figure 3:**
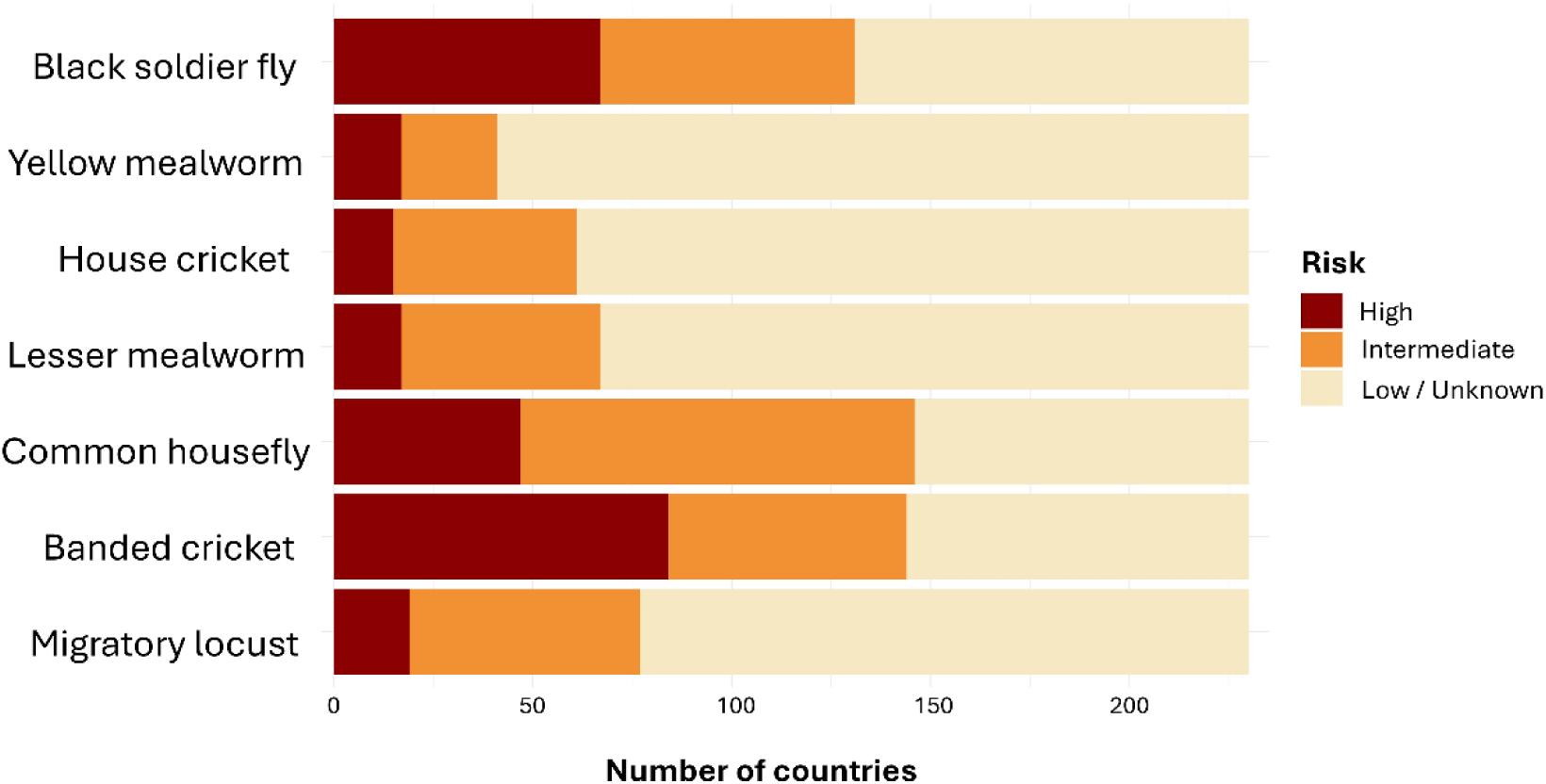
Number of countries in each invasion risk category (low or unknown, intermediate or high) for each of the seven most farmed insect species. Invasion risk categories were defined using quantiles of species occurrence values along the habitat suitability gradient. Habitat suitability values at or below the 5th percentile were classified as low (or unknown) risk, those between the 5th and 25th percentiles as intermediate risk, and those above the 25th percentile as high risk.

Our analysis of risk percentages across IPBES subregions indicates that Central and Western Europe are the most at-risk, followed by Oceania, South Africa, the Caribbean, and South America (**Fig. 4**). The black soldier fly poses the greatest risk in the Caribbean, while the yellow mealworm, house cricket, lesser mealworm, common housefly, and migratory locust are most at risk in Central and Western Europe. The banded cricket is particularly at risk in Oceania, the Caribbean, and Mesoamerica (**Fig. 4**).

**Figure 4:**
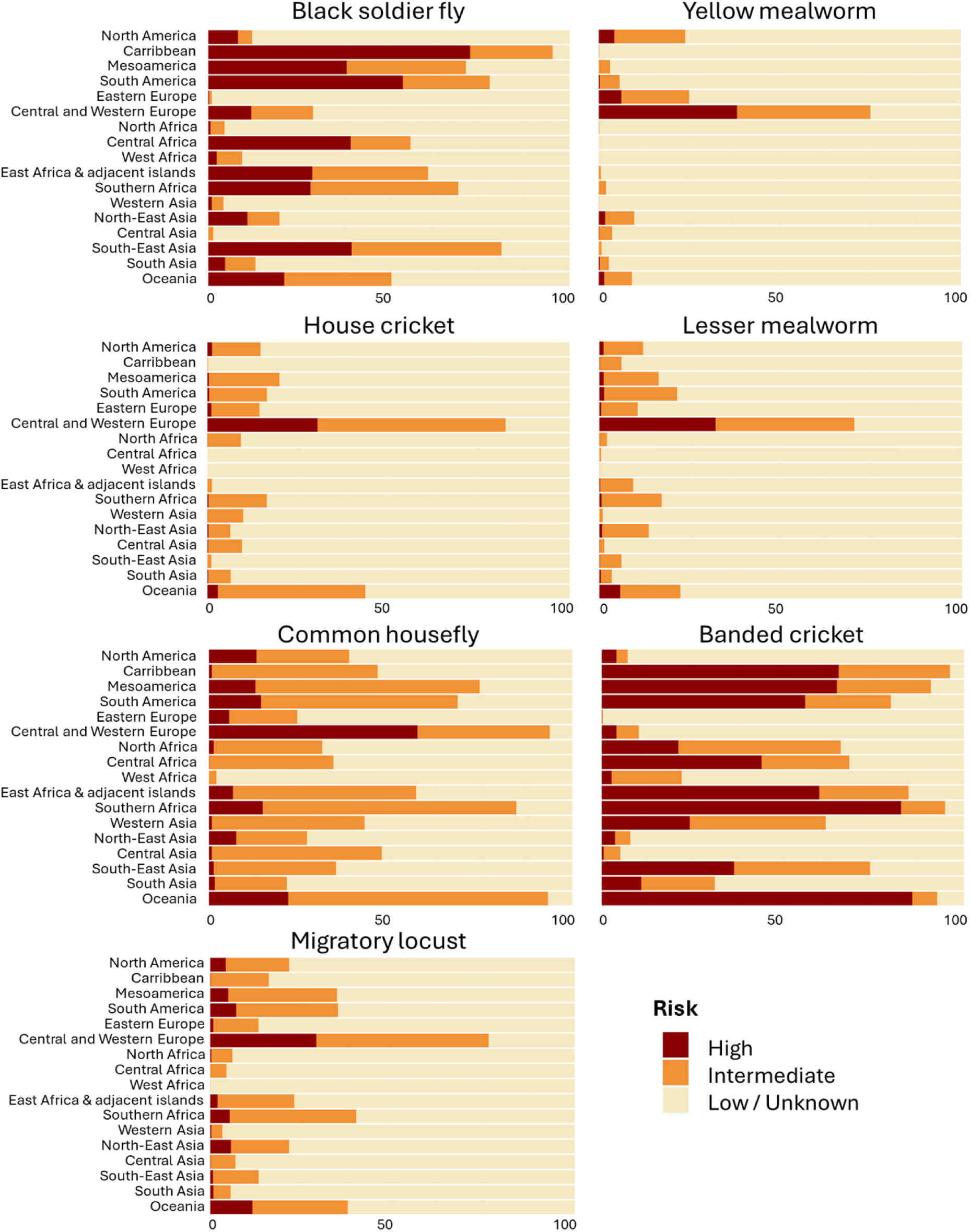
Percentage of invasion risk (low or unknown, intermediate, high) of all IPBES sub-regions for the seven most farm species.

### Risk assessment of insect farming

We sampled a total of 328 farm locations (**Fig. 5,A**) on the basis of our internet searches, categorizing them by taxonomic order since in most cases the exact taxon name could not be retrieved. We found Orthoptera to be the most frequently farmed (102 farms), followed by Diptera (94) and Coleoptera (47) (**Fig. 5,B**), and 21 farms reared multiple species orders. We found that most (80 to 96%) of the farms in each taxonomic category are situated in areas projected to be at intermediate to high invasion risks. This risk was especially prominent for Diptera where 80% of the farms were located in high-risk areas, compared to Orthoptera (55%) and Coleoptera (51%).

**Figure 5:**
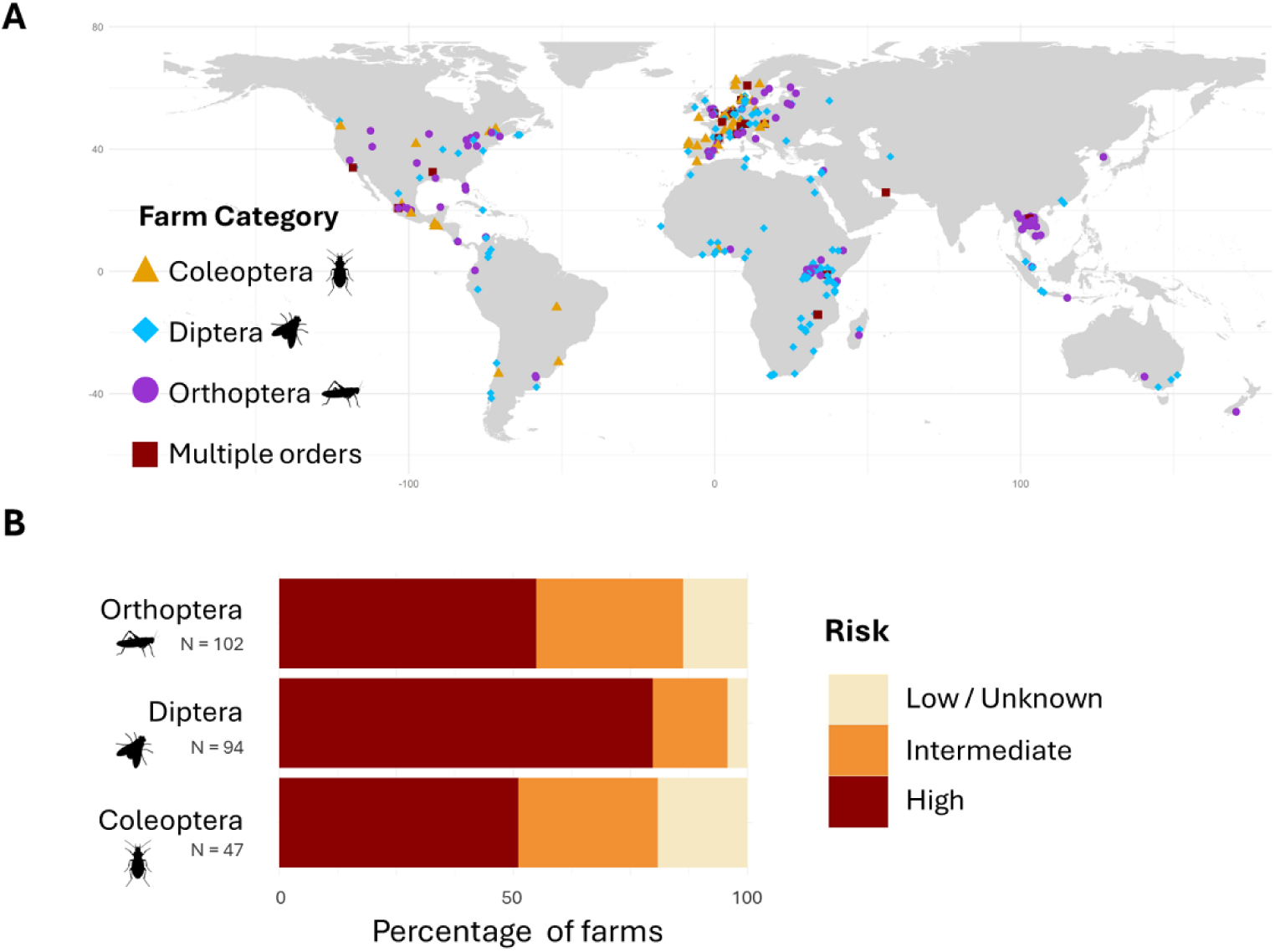
Geographical distribution of insect farms by taxonomic order (Panel **A**) and taxonomic risk assessment of farmed insects (Panel **B**).

## Discussion

Our study demonstrates and quantifies for the first time the biological invasion risks associated with insect farming, addressing concerns raised previously^12,13^. Our results compellingly establish that the large-scale farming of insects poses high invasion risks across the world, given the large extent of intermediate to high suitability areas for the seven most reared insects, and the location of insect farms predominantly in at-risk areas. These results shed light on a major issue that requires attention from the scientific and professional communities working on insect rearing, as well as from policy-makers. If we are to avoid repeating the costliest mistakes of the past^15,28^, we must impulse a concerted effort to prevent the invasion risks posed by this growing industry, or we can expect to pay the cost of inaction^35^. It is very well established that prevention is the best weapon against biological invasions^14^, as eradication programs are the most expensive and often ineffective for insect invasions^36,37^. Our results are crucial tools, enabling strategic species selection and insect farm location in low-risk areas, or the enforcement of maximal biosecurity measures in areas with intermediate to high invasion risks.

### Considering invasive potential: a critical step toward sustainable insect farming

#### Well-established invasion risks for the seven most farmed species

By revealing a substantial and global establishment risk for the seven most commonly farmed insect species, we demonstrated a high potential for biological invasions, with significant socio-economic and ecological consequences. Several of these farmed species are already recognized as pests or invasive, with documented impacts on agriculture, human quality of life, and public health. For instance, the lesser mealworm damages poultry farms^18^, the common housefly affects milk and egg production, while causing sanitary issues^38^, and the migratory locust devastates crops across Africa and Asia^39,40^. Moreover, the common housefly contaminates food and waste^38^ and both the lesser mealworm and common housefly transmit pathogens such as *Salmonella*, *Escherichia coli*, *Shigella*, and *Vibrio cholerae*^41,42^.

These species require particular attention and monitoring due to their high invasive potential, having been selected for traits that enhance establishment risk such as high phenotypic plasticity, generalist habits, disease resistance and rapid life cycles^43–45^. High phenotypic plasticity bolsters tolerance to variable environmental conditions experienced during the invasion process^46^. Combined with high fecundity, these traits allow these species to thrive in a wide range of environments and habitats. With such high adaptability, these species could further exacerbate their invasive potential under climate change, which impose rapid and widespread range expansions^45,47,48^.

#### Invasion risks beyond the seven most farmed species

We provided biological invasion risk assessments for seven species, yet many more species are farmed worldwide, including some with well-known histories of invasion such as the red palm weevil (*Rhynchophorus ferrugineus*)^43,49^. Therefore, biological invasion risks highlighted in this study are likely to be similar for other already farmed species. Furthermore, as the insect market continues to grow^30,31,50^, we can expect an increase in the number of farmed species, necessitating the consideration of invasion risks for future selected species. New farmed species are likely to be chosen among the 1,600 insect species edible by humans worldwide^51^ . However, some of these species are already invasive; three are among the 100 worst invasive species globally (the asian longhorned beetle (*Anoplophora glabripennis*), the common wasp (*Vespula vulgaris*), and a subspecies of the formosan termite (*Coptotermes formosanus*)^52^) and one is on the list of invasive alien species of Union concern (the asian hornet (*Vespa velutina*)^53^). Therefore, the biological invasion risks of farmed species must be considered for both current and future farmed species. Species-specific risk assessments, like in this study, should guide careful selection by policymakers and industry stakeholders, especially since introduced agricultural species often cause the greatest economic damage^54^.

### Strategies to mitigate the risk of biological invasions

#### Species selection

A key objective of the insect farming industry is to reduce food insecurity, particularly in developing regions^9^. Currently, these regions predominantly farm non-native species (e.g., in aquaculture^55^) and are more at risk and disadvantaged in tackling biological invasions^56,57^. Therefore, selecting appropriate species for farming to minimize invasion risks should be a primary concern in farm implementation. Priority should be accorded to native species, as they inherently pose the lowest risk of invasiveness. Developing regions traditionally consume native insect species harvested from the wild^9,43^. Research efforts should therefore prioritize these already-consumed native species to assess their suitability for industrial applications^58^. By focusing on these species, it may be possible to significantly enhance food security while concurrently mitigating the risks of biological invasions in these areas. A moderate-risk option is to farm species in non-native regions where the invasion risk is projected to be low by SDMs and complemented by local risk assessments. For example, the black soldier fly would be better suited for farming in Europe than the common housefly, which in turn would be more appropriate to farm in southeast Asia, where the black soldier fly poses a high invasion risk. The highest-risk category is farming species in non-native areas favorable for their establishment, which must be avoided in new farm implementations to prevent the unavoidable damages and management cost caused by invasions^18,35^.

#### Biosecurity measures

Even though the number of farms here was much underestimated - with Thailand alone reported to have over 20,000 insect farms^59^ - the predominant proportion of farms located in intermediate to high-risk areas underscores the urgent need for stringent biosecurity measures to prevent new biological invasions. Our findings can be expanded to estimate the risk category for all farms rearing these seven species, enabling the implementation of appropriate biosecurity measures.

Regardless of farm size, robust biosecurity measures should be mandatory for farming non-native species. Even for native species, particularly in large-scale industries, stringent biosecurity measures should be enforced to mitigate the ecological impact of mass introductions^60,61^. Effective biosecurity measures are not limited to preventing escapes, but also include conducting regular monitoring of surrounding regions.

#### Policy implications

While challenges in the production and consumption of insects for humans have been raised^62^, and the necessity for biosecurity to ensure sustainable food production and safety acknowledged^63^, measures addressing the biological invasion risk of this industry are largely nonexistent^13^. More broadly, despite the international community’s increasing desire to act on biological invasions, there is a global lack of policies, which are especially lacking in developing regions where the risk and impact of biological invasions are most significant^14,54,64^. By identifying areas at risk of becoming invasion hotspots, our results provide a framework to assist policymakers and industry stakeholders in regulating the farming of high-risk species or strategically locating insect farms in zones with minimal ecological risk.

### Methodological insights and limitations

To optimally serve as a basis for assisting policymakers and industry stakeholders, our results however require further local risk assessment. Here, they provide an overview of potential risks, with limitations inherent to global models. Our comparative analysis of various modeling approaches highlights the critical importance of informed decision-making at each stage of SDMs, as these choices can significantly influence model outcomes and their interpretation^33^. Aligning methodological choices with research objectives presents a notable challenge. For example, two studies predicting future distributions of the migratory locust and the black soldier fly under climate change scenarios employed divergent modeling strategies. One study used six algorithms for model fitting^65^, while the other relied on a single algorithm^66^, leading to substantial differences in their predictions. Two key strengths of our approach is the critical evaluation of models with expert knowledge^67,68^ and the method used to generate pseudo-absences^69,70^, which enhance the ecological realism of our predictions. Specifically, our pseudo-absence protocol aims at reducing effects of sparse and biased occurrence data^71^, a prevalent issue in insect distribution data^72^. Compared to more standard approaches which rely on sampling background points, this protocol generally produces larger potential ranges, which is a desirable characteristic in biological invasions where the aim is to assess a realistic risk as largely as possible^73^. However, the generalisability of this approach has not yet been demonstrated and would benefit from further applications and comparisons in the future. Despite providing an upper boundary of the global invasion risk, this method may still underestimate this risk in certain regions, such as for example in India, where the black soldier fly and the housefly have numerous occurrences despite the country being classified as at low risk by our models. This example is illustrative of the limits of our models which may not fully capture the full array of drivers of species distributions.

On the one hand, SDMs are correlative approaches which make it difficult to include drivers related to dispersal or biotic interactions, especially for insects which have larger knowledge gaps than vertebrates^74^. On the other hand, knowledge shortfalls on the distribution data *per se*^72^ may result in the lack of identification of important drivers. For example, the migratory locust inhabits difficult-to-access regions such as the central Niger delta and the Blue Nile region of Sudan^75,76^, leading to an absence of occurrence data to feed our model and, consequently, an underestimated establishment risk in similar areas. Additionally, species can be challenging to identify due to phenotypic similarities with other species, which is the case of the house cricket^77^. Consequently, our models are indicative of the areas suitable given the climatic and socio-economic drivers included, which we assume to be the most important at a large spatial scale.

Our study represents one of the first opportunities to be proactive and anticipate major environmental and socio-economic risks posed by a fast-growing industry. Our global-scale results are sufficient to alert policymakers to the significant risk associated with insect farming and provide a strong basis for policy decisions and optimal strategic planning. This study also represents a first concrete step in this research area and underscores the need for further research to help the food industry achieve its objective of balancing the advancement of sustainable protein production with the preservation of native biodiversity.

## Methods

The common housefly, the black soldier fly, the domestic cricket, the lesser mealworm and the yellow mealworm have become cosmopolitan as a result of long-term human association and transportation, and their native regions remain debated^78–80^. Due to this uncertainty, we chose to not exclude any potential native regions from our SDM projection maps, ensuring that all possible spatial risks are accounted for.

### Data preparation

#### Species occurrences

We compiled species occurrence records from the Global Biodiversity Information Facility (GBIF, https://www.gbif.org/) using the “rgbif” package^81^. To improve the accuracy of the occurrences, we selected data from 1980 to 2024, a period benefiting from the advent of the Global Positioning System and advancements in data digitization. We downloaded species occurrences ranging from 143 (for the field cricket) to 9953 (for the black soldier fly) (**Supplementary Table S1**).

#### Environmental and socioeconomic variables

A key factor in assessing the biological relevance of distribution models is to select variables according to the taxonomic group ecology^67^. Although species in our study can tolerate a wide range of climatic conditions, their development is particularly sensitive to temperature and humidity^45,82–84^. We selected these key environmental variables from Climatologies at High Resolution for the Earth’s Land Surface Area (CHELSA, 1 km pixel resolution; chelsa-climate.org^85^). These species are generalist feeders, primarily phytophagous, with some displaying an omnivorous diet^45,78,79,86–88^. They are also generally closely linked to human activity, feeding on food waste or crops^38,40,78^. We therefore have considered three human-related variables including *Croplands* (1 km pixel resolution^89^), *Human population* density (1 km pixel resolution^90^) and *Human footprint* (1 kilometer pixel resolution^91^). After download, we harmonized each climatic and socioeconomic variable onto a standard of 5 kilometer pixel resolution. We finally excluded all variable conditions outside land using Natural Earth data (https://www.naturalearthdata.com/downloads/) and cropped the extent, considering a maximum latitude of 75° and a minimum latitude of -60°.

### Preliminary variable selection

We assessed variable correlations using the Pearson correlation coefficient (r > 0.6^92^) to identify groups of correlated variables. Within each group, we evaluated pairwise correlations to eliminate redundant variables, prioritizing those with higher biological relevance to insect ecology (**Supplementary Fig. S7**). This process resulted in the selection of four climatic variables: mean daily maximum air temperature of the warmest month (*bio5*), mean daily minimum air temperature of the coldest month (*bio6*), minimum surface relative humidity (*hurs_min*), along with net primary productivity (*npp*). Additionally, we included the three socioeconomic variables - *croplands*, *human population*, and *human footprint* - analyzing them separately to develop models that combine climatic variables with one socioeconomic variable.

### Environmental filtration

Because SDMs rely on species occurrence data, they require absence data for model calibration. However, absence data is unavailable for our farmed species and must be inferred. Traditional approaches generate absences randomly or with geographic biases, potentially compromising model accuracy^93,94^. An alternative method involves estimating a species’ suitable environmental conditions using occurrence data and ecological characteristics by representing them in a convex hypervolume (called a convex hull). Pseudo-absences are then generated outside this convex hull. Unlike conventional geographic filtering, this environmentally driven approach has been shown to significantly enhance model performance^69,71^.

#### Environmental space

We created the environmental space by extracting values for each of the five variables and projecting them into an environmental grid. This grid represents a multidimensional space where each axis corresponds to one of the environmental variables, divided into 80 equal intervals.

#### Occurrence filtration

We filtered species occurrences by first removing those with missing variable values. Expert scientists then analyzed the remaining occurrences, which were subsequently projected into the environmental space. This process ensured that only unique and realistic species occurrences, along with their respective variable values, were retained in the environmental space (**Supplementary Table S1**).

### Pseudo absence generation

We generated pseudo-absences by first creating a convex hypervolume (convex hull) that represents the species’ environmental conditions, based on predefined environmental variables and species occurrences. To eliminate probable false occurrences, we removed extreme values by discarding the 0.5% lowest and highest values for each variable at the occurrence points. We hypothesized that environmental conditions within the convex hull are favorable for species survival. Therefore, we generated pseudo-absence points randomly outside the convex hull, ensuring they did not overlap with the extreme occurrences (1% per variable) excluded from its construction. We created five sets of pseudo-absences, each equal in number to the occurrence points, resulting in a total number of pseudo-absences five times greater than the occurrence data. (**Supplementary Fig. S8**).

As we have no prior assumptions about the shape of the relationships between species and environmental variables, we used three machine learning and two statistical models known for their strong predictive performance in presence only situations^33,95^. These models are (i) Maxnet (ii) extreme gradient boosting (XGBOOST), (iii) Random Forest downsampled (RF), (iv) General Additive Models (GAM) and (v) General Linear Models (GLM).

These analyses and the following ones were conducted using the Biomod2 package and its associated functions^96^. To ensure optimal performance, we optimized the parameters for RF and XGBOOST for each species (see model parameters in **Supplementary Table S2**).

### Model selection and evaluation

We estimated the probability of species occurrence along environmental gradients using our models, aiming for realistic and appropriately complex response curves to avoid overfitting (see response curves in **Supplementary S3**^68,71,97^). To achieve this, we randomly divided each dataset into five subsets (or cross-validation folds), each containing an equal number of pseudo-absences and occurrences. Each fold was used for evaluation, while the remaining four were used for model calibration, resulting in four training iterations per run. We assessed model performance using the Boyce Index, which ranges from -1 (indicating model predictions not consistent with occurrence distributions) to 1 (indicating model predictions consistent with occurrence distributions), and retained models scoring above 0.7. Expert scientists then analyzed the realism of the response curves. Finally, we estimated variable importance by measuring the performance loss when each variable was removed (**Supplementary S3**).

### Species distribution projections

#### Ensemble model

To predict the global risks of establishment, we projected each individually calibrated model and calculated an ensemble model by calculating the average of individual projections. For each species, these ensemble models represent our estimation of the risk of species establishment (see maps in **Supplementary S4**). Finally, we calculated the standard deviation among the models to quantify the uncertainty of the modeling process (see maps in **Supplementary S4**).

#### Invasion risk

We categorized the invasion risk by analyzing the distribution of occurrences along the habitat suitability gradient generated by the ensemble model, using a method based on quantiles of occurrences at the 5th and 25th percentiles^98^. We classified habitat suitability values at or below the 5th percentile as low (or unknown) risk, those between the 5th and 25th percentiles as intermediate risk, and values above the 25th percentile as high risk (**Supplementary S6**). This classification ensures that each category of invasion risk conveys a similar meaning across species, enabling a comparative assessment of invasion risk. To analyze the geographic distribution of invasion risk, we utilized the regional classification provided by IPBES^99^.

### Insect farm locations

To assess the actual risk of insect farming, we examined the location of insect farms within risk invasion categories. To do so, we gathered insect farm coordinates from various online sources, including Google Maps searches in multiple languages (primarily French, English, Spanish), and extracted data from scientific studies using QGIS^31,59^. We collected coordinates of 328 farms located across all continents (**Supplementary Table S3**). Due to the limited availability of precise scientific names, we categorized insect farms by the taxonomic order of the species they farm: Orthoptera (house cricket, banded cricket, field cricket, migratory locust), Diptera (black soldier fly, domestic housefly), and Coleoptera (yellow mealworm and lesser mealworm). Our objective with this dataset was not exhaustivity, nor precise representativity, but rather obtaining a global idea of the quantity and repartition of the localisation of the facilities of different species of farmed insects.

## Acknowledgments

This study received support from support from the AXA Research Fund Chair for Biological Invasions at the University of Paris Saclay.

## Author contributions

Conceptualization: EM, BL, FC, QL

Methodology: EM, BL

Analysis: EM

Writing—original draft : EM

Review : EM, BL, FC, POM, QL

## Notes

https://github.com/Elena-Manfrini/SDM_Insect-farming/tree/v1.0

### Competing Interest Statement

The authors have declared no competing interest.

